# Pre-existing systemic and nasal antibodies against avian H5 influenza A viruses vary according to childhood imprinting

**DOI:** 10.64898/2026.05.08.723737

**Authors:** Peta Edler, Kevin J. Selva, Ellie Reilly, Malet Aban, Ian G. Barr, Jennifer A. Juno, Adam K. Wheatley, Amy W. Chung, Stephen J. Kent, David J. Price, Marios Koutsakos

## Abstract

Avian influenza A viruses (IAV) pose a constant pandemic threat, with the recent 2.3.4.4b clade of the H5 subtype causing high pathogenicity and spreading across animal species and geographic locations. Understanding human pre-existing immunity to avian H5 IAV can inform on population susceptibility, a critical aspect of pandemic preparedness. To that end, we analysed the IAV HA-specific antibodies across individuals born between 1928-1999 with different early life exposures to IAV subtypes. Individuals born prior to 1957 had the highest pre-existing serum antibodies to group 1 HA antigens, including the 2.3.4.4b H5 and a group 1 HA stem antigen. These birth-year-specific patterns were not reflected in the limited pre-existing serum neutralising antibodies detectable against a 2.3.4.4b H5 IAV or in H5-specific memory B cell populations. They were however evident in pre-existing nasal IgG and IgA titres to H5, which were greater in individuals born prior to 1957. Our findings demonstrate that the immunological biases afforded by early life exposure extend to antibodies detected in the nasal mucosa, the site of IAV replication.

**Importance:** Understating pre-existing immunity to influenza A viruses of pandemic potential is an important aspect of pandemic preparedness. This includes an understanding the heterogeneity of pre-existing immunity across the population. Here, we demonstrate that pre-existing antibodies to H5 IAV vary according to year of birth and childhood imprinting. We demonstrate that this is the case for both systemic and nasal antibodies, highlighting the importance of understanding pre-existing mucosal immunity at the sites of influenza virus replication.

## Introduction

Highly pathogenic avian influenza (HPAI) viruses of the H5 subtype currently represent an evolving threat to both animals and humans. Since 2020, clade 2.3.4.4b H5N1 and related H5Nx viruses have caused unprecedented outbreaks in wild birds and poultry globally(1). The sustained infections of diverse mammalian hosts, including wild and farmed mammals, raise concerns for increased opportunities for viral adaptation in mammalian hosts. Human infections with contemporary H5N1 viruses remain sporadic and are typically associated with exposure to infected animals, with no evidence of sustained human-to-human transmission(2). Nonetheless, the wide geographic spread and repeated mammalian spillovers underscore the need for intensified surveillance, continued development of pre-pandemic H5 vaccines and therapeutics, and an increased understanding of pre-existing population immunity.

Pre-existing immunity is an important determinant of influenza pandemic outcomes, influencing both individual susceptibility and population-level burden(3, 4). Such pre-existing immunity can be provided by cross-reactive antibodies elicited by prior exposures to seasonal influenza A viruses (IAV). In the context of novel IAV subtypes the main target of such cross-reactive antibodies is typically the highly conserved stem domain of the HA, although other conserved antibody targets on the HA and NA exist(5, 6). Since antibody responses to the HA stem have been associated with protection from seasonal influenza virus symptomatic infection(7, 8), cross-reactive antibodies to the HA stem of avian influenza viruses may also contribute to a degree of protection. Therefore, understanding the degree of such pre-existing immunity across the population can inform overall population risk. However, population immunity is not homogeneous, and it is important to understand variability to identify high-risk groups. This is further supported by the epidemiology of zoonotic infections with H5 IAV, whereby younger adults have been more susceptible to severe disease and fatal infection than older individuals(9).

This differential susceptibility between younger and older adults has been associated with different early life exposures to IAVs from different HA groups(9). Specifically, those born prior to 1957 were first exposed to an H1N1, a group 1 HA, and thus show reduced susceptibility to avian H5 IAV, also a group 1 HA. A similar pattern has also been observed for those born between 1957-1968, who were most likely first exposed to an H2N2, a group 1 HA. Conversely, those born between 1968-1976 were first exposed to an H3N2 subtype, a group 2 HA, and thus show reduced susceptibility to avian H7 IAV, also a group 2 HA. It has been proposed this is a result of immunological imprinting, whereby first exposures to a specific virus result in a long-term immunological bias that shapes subsequent susceptibility(9–11). Recent serological studies have indeed indicated that pre-existing immunity to H5 IAV in individuals with no prior A(H5) exposure) varies by year of birth(12, 13) and likely explains this differential susceptibility. These studies have focused on the analyses of serum antibodies, however, whether differences in pre-existing immunity are also reflected in memory B cells (MBCs) or mucosal antibodies is unclear.

Here we demonstrate that (i) pre-existing serum antibodies to group 1, but not group 2, IAV HAs vary by year of birth and thus according to childhood imprinting; (ii) these differences are not reflected in pre-existing serum neutralising antibodies against H5 IAV or H5-specific memory B cell (MBC) populations; (iii) these differences are reflected in pre-existing nasal IgG and IgA antibodies to H5. Our findings demonstrate that the immunological biases afforded by early life exposure extend to antibodies detected in the nasal mucosa, the site of IAV replication.

## Results

### Pre-existing serum binding antibodies against avian H5 IAV vary by birth year

To understand how antibodies to seasonal and avian IAV vary by childhood imprinting we analyzed serum samples from 144 individuals born between 1928 and 1999 and sampled between 2020-2023 (Figure 1A). These individuals represent 4 birth cohorts with different early life exposure to IAV subtypes, which can be inferred from the historical circulation patterns of IAV(9, 12): individuals born prior to 1957 (1928-1956 in our cohort) were most likely imprinted with the group 1 H1 subtype; individuals born between 1957-1967 were most likely imprinted with the group 1 H2 subtype; individuals born between 1968-1976 were most likely imprinted with the group 2 H3 subtype; and individuals born after 1977 (1977-1999 in our cohort) may have been imprinted with either the group 1 H1 subtype or the group 2 H3 subtype. Using these samples, we assessed how binding serum IgG antibodies of different IAV HA specificities vary by year of birth. As these individuals do not have known exposure histories to avian IAV, the detection of antibodies specific for avian IAV HAs represents pre-existing cross-reactive immunity to these antigens.

**Figure 1.**
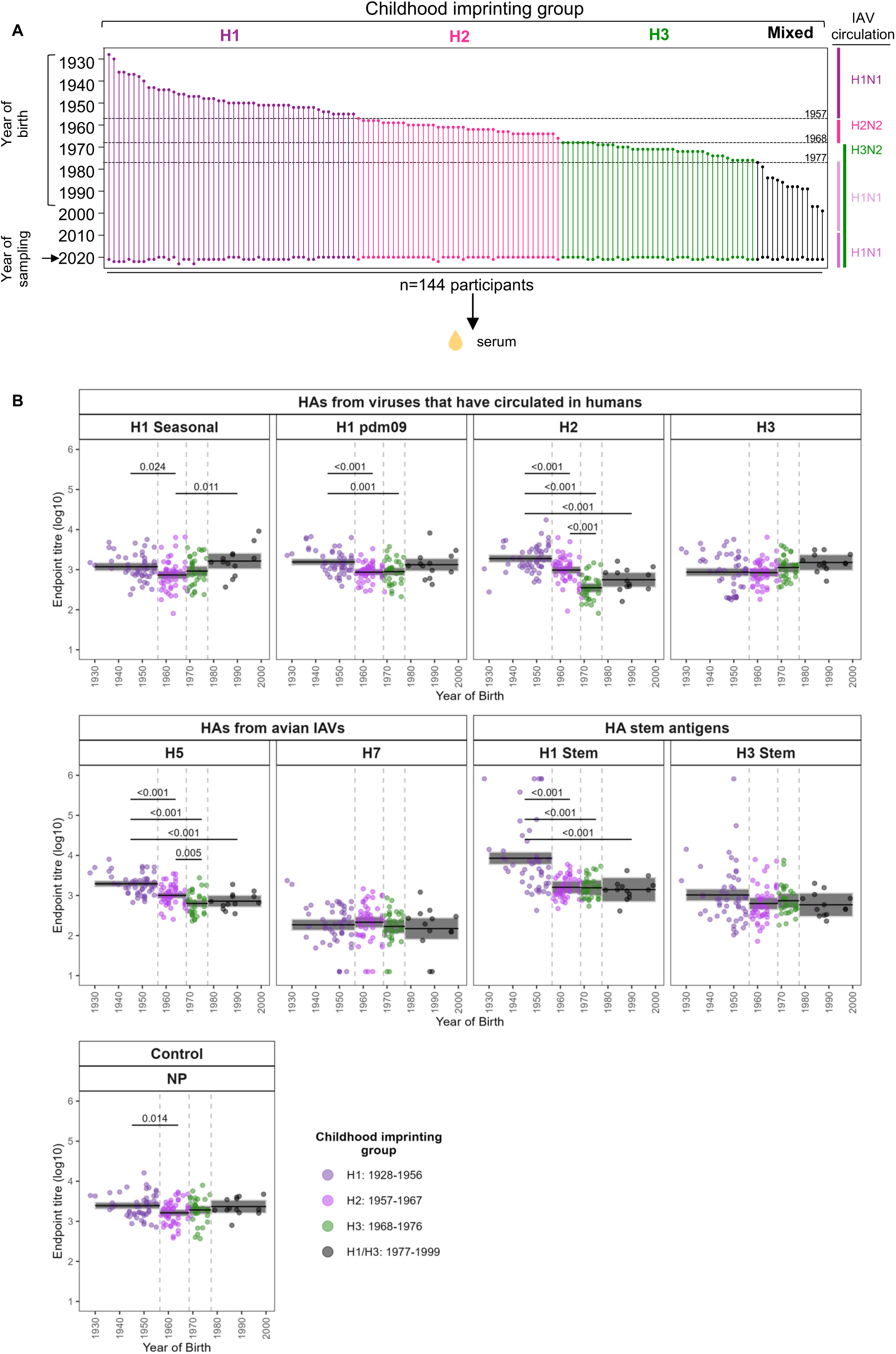
Pre-existing serum IgG antibodies against avian H5 IAV vary by birth year. **(A)** Cohort description. Each vertical line represents a donor with the top circle representing their year of birth (1928-1999) and the bottom circle representing their year of sampling (2020-2023). Horizontal lines indicate the year new IAV subtypes were introduced, and the subtype circulation patterns are indicated on the right. The inferred childhood imprinting groups are indicated on the top. **(B)** Antibodies against HAs from Influenza A viruses (IAVs) and stem antigens by year of birth. Each panel represents a different IAV, stem antigen or control. Childhood imprinting groups are represented by the coloured points. The black lines represent the mean for each imprinting group, and the grey bars represent the 95% confidence interval of the mean. Between-group comparisons, which resulted in an adjusted p-value (false discovery rate) greater than 0.05, have been shown.

These imprinting patterns were reflected in antibodies against the HA protein for IAVs that had circulated in humans (H1, H2 and H3) (Figure 1B)(Table 1). Those born prior to 1957 and after 1977, who were likely imprinted with group 1 H1 subtypes, had the highest antibodies against HAs from H1 subtypes (A/Sydney/5/2021 H1pdm09 and A/Solomon Islands/03/2006 seasonal H1). Antibodies against the H2 subtype (A/Canada/720/2005) were the highest in those born between 1928-1956, followed by those born between 1957-1968, when the H2 subtype circulated. In contrast to these antibodies against group 1 HAs, antibodies against H3 (A/Darwin/6/2021), a group 2 HA, were not different between birth cohorts. Similar patterns were observed for antibodies against group 1 and 2 HAs from avian IAVs that have not circulated in humans (H5 and H7), with a birth year bias observed against HA proteins from the H5 subtype (A/Fujian-Sanyuan/21099/2017), but not from the H7 subtype (A/Shanghai/02/2013). Specifically, those most likely imprinted with a group 1 HA (H1 or H2) had higher H5 titres than other birth cohorts, while antibodies to H7 were uniformly low across birth cohorts. These patterns were also consistent when considering antibodies against the HA stem domain from group 1 (H1 stem) and group 2 (H3 stem) HAs. Those most likely imprinted with H1 had higher group 1 stem titres than other birth cohorts, while antibodies to group 2 stem were uniform across birth cohorts. Overall, these data suggest that serum IgG antibodies against group 1 but not group 2 IAV HAs vary according to childhood imprinting.

**Table 1:** Mean and 95% confidence intervals for antibodies against HAs from influenza A viruses (IAVs) and stem antigens by childhood imprinting group.

|  | **Mean (95% CI) antibodies per childhood imprinting group** |  |  |  |
| --- | --- | --- | --- | --- |
| **HAs from viruses that have circulated in humans** | **H1: 1928-1956** | **H2: 1957-1967** | **H3: 1968-1976** | **Mix: 1977-1999** |
| H1 seasonal | 3.07 (2.97, 3.17) | 2.87 (2.77, 2.97) | 2.96 (2.84, 3.09) | 3.21 (3.02, 3.41) |
| H1 pdm09 | 3.19 (3.11, 3.27) | 2.94 (2.86, 3.02) | 2.95 (2.85, 3.05) | 3.12 (2.97, 3.28) |
| H2 | 3.28 (3.18, 3.37) | 2.99 (2.90, 3.08) | 2.55 (2.43, 2.66) | 2.75 (2.57, 2.93) |
| H3 | 2.94 (2.84, 3.04) | 2.92 (2.82, 3.03) | 3.05 (2.93, 3.18) | 3.18 (2.98, 3.37) |
| **HAs from avian IAVs** |  |  |  |  |
| H5 | 3.29 (3.22, 3.37) | 3.00 (2.93, 3.08) | 2.80 (2.71, 2.89) | 2.86 (2.71, 3.00) |
| H7 | 2.27 (2.13, 2.40) | 2.33 (2.19, 2.47) | 2.23 (2.06, 2.40) | 2.17 (1.91, 2.44) |
| **HA stem antigens** |  |  |  |  |
| H1 stem | 3.93 (3.78, 4.08) | 3.20 (3.05, 3.36) | 3.19 (3.00, 3.38) | 3.15 (2.84, 3.45) |
| H3 stem | 3.01 (2.86, 3.16) | 2.80 (2.65, 2.95) | 2.87 (2.68, 3.06) | 2.77 (2.48, 3.06) |
| **Control** |  |  |  |  |
| NP | 3.39 (3.31, 3.47) | 3.21 (3.13, 3.29) | 3.28 (3.18, 3.39) | 3.37 (3.21, 3.53) |

As our samples were primarily collected between 2020-2023 during which there was little to no circulation of influenza viruses in Australia as a result of the COVID-19 pandemic(14, 15), we considered it unlikely that these differences are a result of differential recent exposure to seasonal IAVs, which may boost IgG titers to H5(16, 17). To further rule out this possibility, we measured binding serum IgG antibodies against the IAV nucleoprotein (NP), which should be boosted by recent infection, but is highly conserved between subtypes and therefore should not vary by childhood imprinting. Indeed, we found that titres to the IAV NP were overall similar across birth cohorts (Figure 1) (Table 1). In addition, only 13% of our samples were collected during the influenza vaccine season (mid-April until August), with 29% of the samples collected outside of that time, and 58% of samples collected prior to seasonal influenza vaccination as baseline samples of a vaccine cohort. We therefore consider it unlikely that the differences between birth cohorts in serum HA titres reflect differences in recent vaccination or infection history.

To further probe these biases in pre-existing cross-reactive antibodies to H5, we considered the pairwise correlations of antibody measurements (Figure 2). Overall, we observed positive correlations between titres against group 1 HA antigens (H1 seasonal, H1 pandemic, H2 and H5). Antibodies against HA from H5 subtype were correlated with antibodies against H2 and H1 pandemic antigens (correlation co-efficient 0.66 [95% CI: 0.56, 0.74] and 0.60 [0.49, 0.7] respectively). We also observed correlations between antibodies against H7 and antibodies against H3, although the birth year biases for these antibodies were not as evident as those for group 1 HAs (Figure 1B). Antibodies against H5 were also positively correlated with antibodies against the group 1 HA stem antigen (0.54 [0.41, 0.64]) and less so with the group 2 H3 stem (0.26 [0.10, 0.40]). This was less evident for antibodies against HAs from both the H1 and H2 subtypes (seasonal: 0.22 [95% CI: 0.06, 0.37]; pdm09: 0.34 [0.18, 0.47]) and H2 subtype (0.24 [0.08, 0.39]) (Figure 2). We also observed correlations between antibodies against H7 and antibodies against the group 2 H3 stem (0.32 [0.17, 0.46]) and less so with the group 1 H1 stem (0.17 [0.01, 0.33]). Overall, these observations support the idea of HA group-level imprinting that results in birth-cohort-specific cross-reactivity against H5, likely via the HA stem.

**Figure 2.**
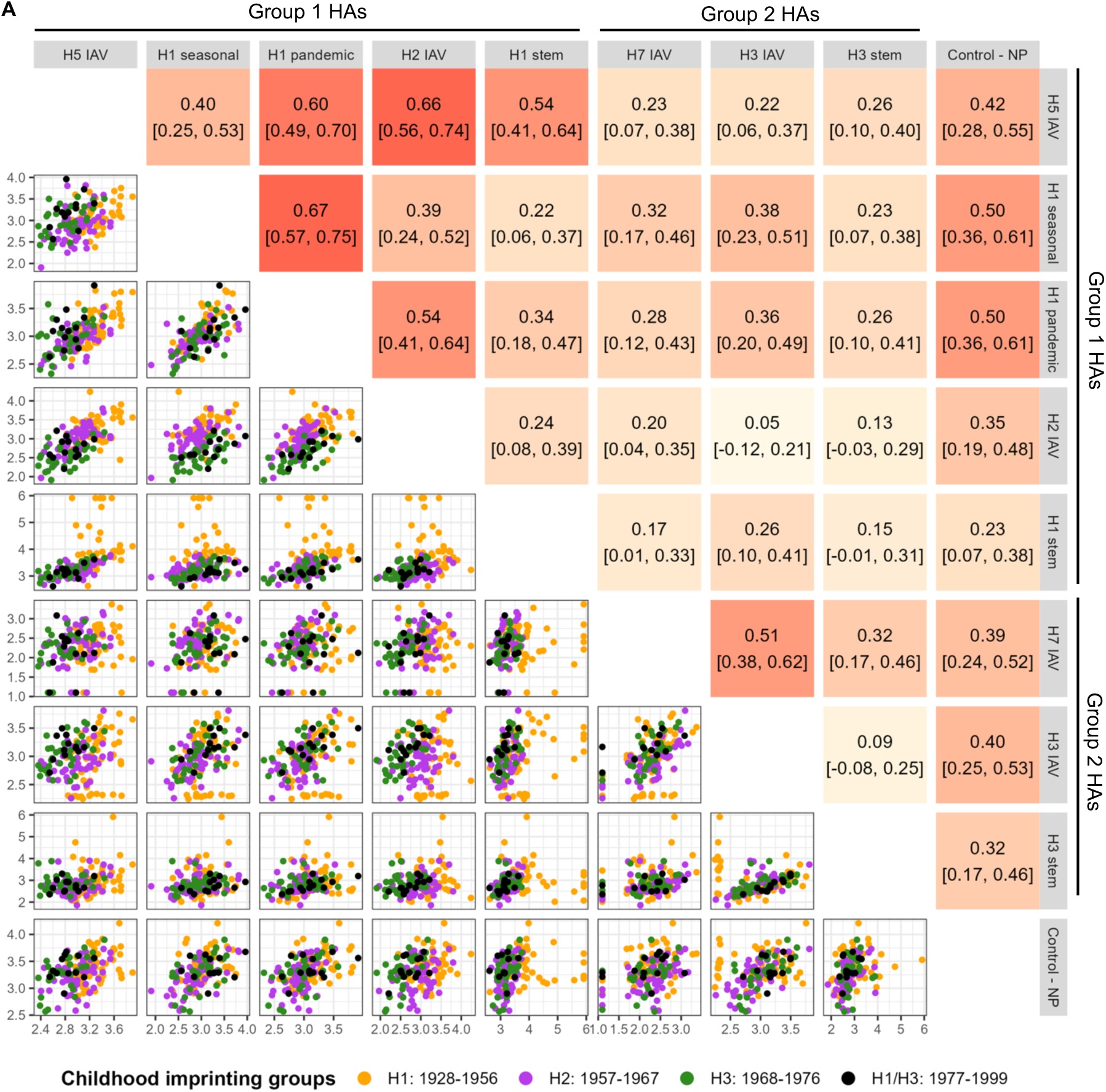
Correlations between antibodies to IAV antigens. Upper triangle: Pearson pairwise correlations with 95% confidence intervals (CIs) for antibody measurements against HA proteins from different IAV subtypes – darker colours squares indicate stronger correlations. Lower triangle: pairwise scatter plots of all HA proteins from different IAV subtypes – point colour corresponds to imprinting group.

### Limited pre-existing serum neutralizing antibodies against avian H5 IAV

To further probe the degree of pre-existing immunity to avian H5 IAVs, we recruited a second cohort of 39 individuals born between 1936-1971, representing the H1-imprinting individuals, H2-imprinting individuals and H3-imprinting individuals (Figure 3A). We collected serum, peripheral blood mononuclear cells and nasal fluid to assess pre-existing antibodies in the circulation and nasal mucosa as well as pre-existing memory B cells (MBCs) to avian H5. Analysis of serum IgG antibodies against H5 (A/Fujian-Sanyuan/21099/2017), a group 1 HA stem and the control NP antigen, reinforced the patterns observed with the first cohort: those imprinted with a group 1 HA in childhood had the highest antibodies against H5 and the group 1 HA stem, while antibodies against NP were similar across birth cohorts (Figure 3B). To understand if these pre-existing antibodies to H5, presumably due to cross-reactivity with the HA stem domain, result in virus neutralization, we performed a live virus microneutralization assays with A/Astrakhan/3212/2020, an H5N8 isolate from the 2.3.4.4b cluster that has been selected as a vaccine candidate strain. Neutralisation activity was only detectable in 4/13 individuals from the H1-imprinted group and 1/13 from the H2-imprinted group (Figure 3C). Even in these individuals, neutralisation titres were low (between 10-20), suggesting limited pre-existing serum neutralizing antibodies against avian H5 IAV.

**Figure 3.**
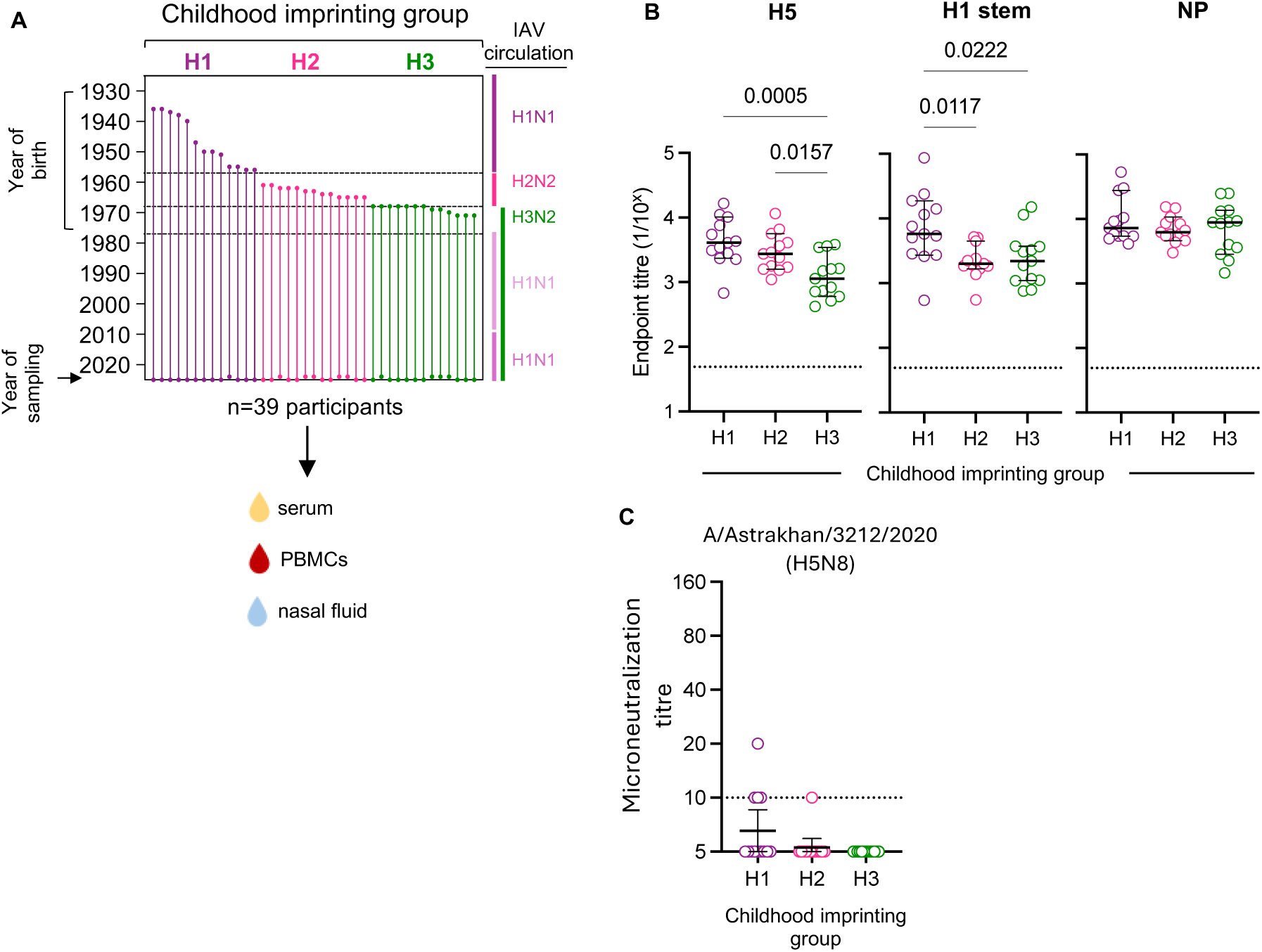
Limited pre-existing serum neutralizing antibodies against avian H5 IAV. **(A)** Cohort description. Each vertical line represents a donor with the top circle representing their year of birth (1928-1999) and the bottom circle representing their year of sampling (2024-2025). Horizontal lines indicate the year new IAV subtypes were introduced, and the subtype circulation patterns are indicated on the right. The inferred childhood imprinting groups are indicated on the top. **(B)** Serum IgG antibody titers to different influenza A antigens from groups with different childhood imprinting to IAV (n=13/group). The median and 95% CIs are shown, p-values were calculated using an ANOVA with Tukey’s correction for multiple comparisons – only comparisons with p-values <0.05 are shown. **(C)** Microneutralization titers to A/Astrakhan/3212/2020 (H5N8) in serum from groups with different childhood imprinting to IAV (n=13/group). The geometric mean titres and 95% CIs are shown, p-values were calculated using the Kruskal-Wallis test with Dunn’s correction for multiple comparisons and were all >0.05. The dotted horizontal lines represent the limit of detection.

### Pre-existing memory B cells against avian H5 IAV do not vary by birth year

We next wanted to determine if individuals with different childhood imprinting histories possess different levels of MBCs against the H5 antigen, as these could possibly impact memory responses following H5 re-exposure in the case of infection or vaccination. We detected MBCs specific for the A/Fujian-Sanyuan/21099/2017 H5 (2.3.4.4b) using flow cytometry (Figure S1) as previously described(18–20). The levels of H5-specific IgG MBCs did not differ between individuals with different childhood imprinting histories (Figure 4A-B). The levels of H5-specific IgG MBCs were modestly correlated (Spearman r<0.5) with the levels of H5-specific serum IgG antibodies but not with the levels of NP-specific serum IgG antibodies (Figure 4C).

**Figure 4.**
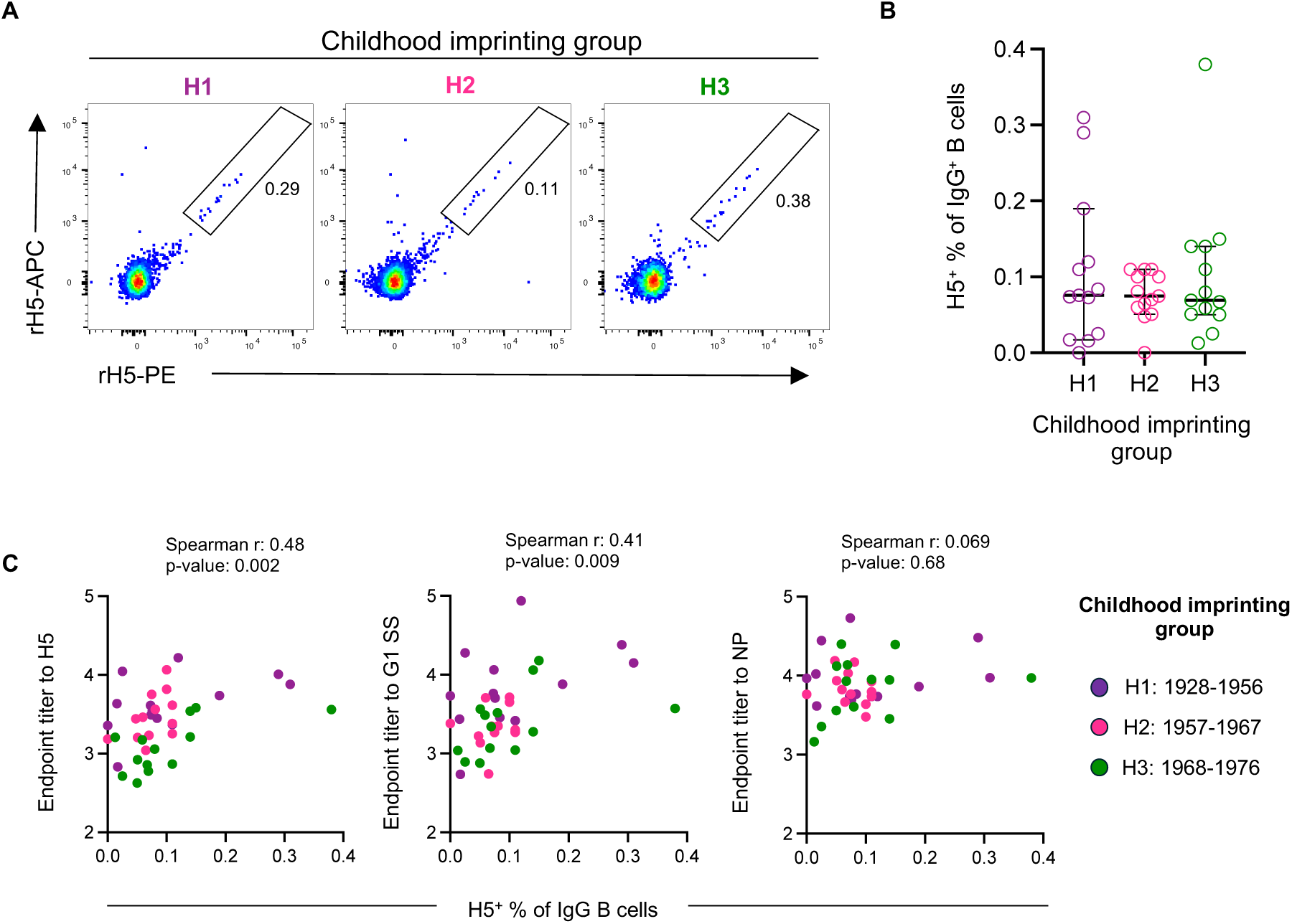
Pre-existing memory B cells against avian H5 IAV do not vary by birth year. **(A)** Representative FACS plots from a donor from each imprinting cohort. A/Fujian-Sanyuan/21099/2017 H5-specific B cells are shown after gating on IgG^+^IgD^-^CD19^+^CD3^-^CD8^-^CD16^-^CD14^-^CD10^-^ live single lymphocytes. **(B)** Frequency of H5-specific cells withing IgG^+^ memory B cells across imprinting cohorts (n=13/group). The median and 95% CIs are shown, p-values were determined with an ANOVA with Tukey’s correction for multiple comparisons and were all >0.05. **(C)** Spearman correlation between antibody titers to H5, group 1 stabilized stem (G1 SS) or NP and the frequency of H5-specific cells (n=39).

### Pre-existing nasal IgG antibodies against avian H5 IAV vary by birth year

Finally, given the importance of mucosal immunity in protection, we wanted to assess if the levels of pre-existing antibodies in the nasal mucosa also vary according to childhood imprinting. We used a multiplex bead assay to measure IgG and IgA antibodies in nasal fluid (Figure S2A). We assessed IgG and IgA antibodies specific for H5 (A/Fujian-Sanyuan/21099/2017); antibodies specific to IAV NP and tetanus toxoid as control antigens that should not vary by year of birth; as well as the total IgG and IgA detected in each sample (Figure S2B). Analysis of serum using the multiplex bead assay with the H5 antigen confirmed the higher levels of H5-specific IgG and IgA in serum of individuals imprinted with H1, followed by those imprinted with H2 (Figure S3A). Antibody titers to the other antigens did not vary by childhood imprinting, as expected. The H5-specific and NP-specific IgG antibody levels detected by multiplex bead assay were positively correlated with the titers determined by ELISA (Figure S3B).

For nasal fluid samples, we considered the antigen-specific MFI normalized to the total IgG or IgA detected in each nasal sample (Figure 5), to account for differences in sampling of the nasal mucosa(21–23). Individuals born prior to 1957 had higher nasal H5-specific IgG (median 0.32, [95% CIs: 0.19, 0.7]) than the other two birth cohorts (0.15 [0.09, 0.26], for H2 group, 0.12 [0.067, 0.24] for H3 group) and higher levels of H5-specific IgA (0.04 [0.017, 0.19] for H1 group, 0.031 [0.014, 0.053] for H2 group, 0.015 [0.005, 0.03] for H3 group). The levels of NP-specific and tetanus-toxoid specific IgG and IgA were similar across the different birth cohorts. The same trends were observed in the raw MFI data (not normalized to total IgG/IgA). The H5-specific IgG, but not IgA, antibody levels detected in nasal fluid were highly correlated to the levels detected in serum suggesting the antibodies detected in nasal fluid are likely derived from transcytosed serum IgG. Overall, our data demonstrate that both pre-existing serum and mucosal antibodies specific for avian H5 IAV vary according to childhood imprinting.

**Figure 5.**
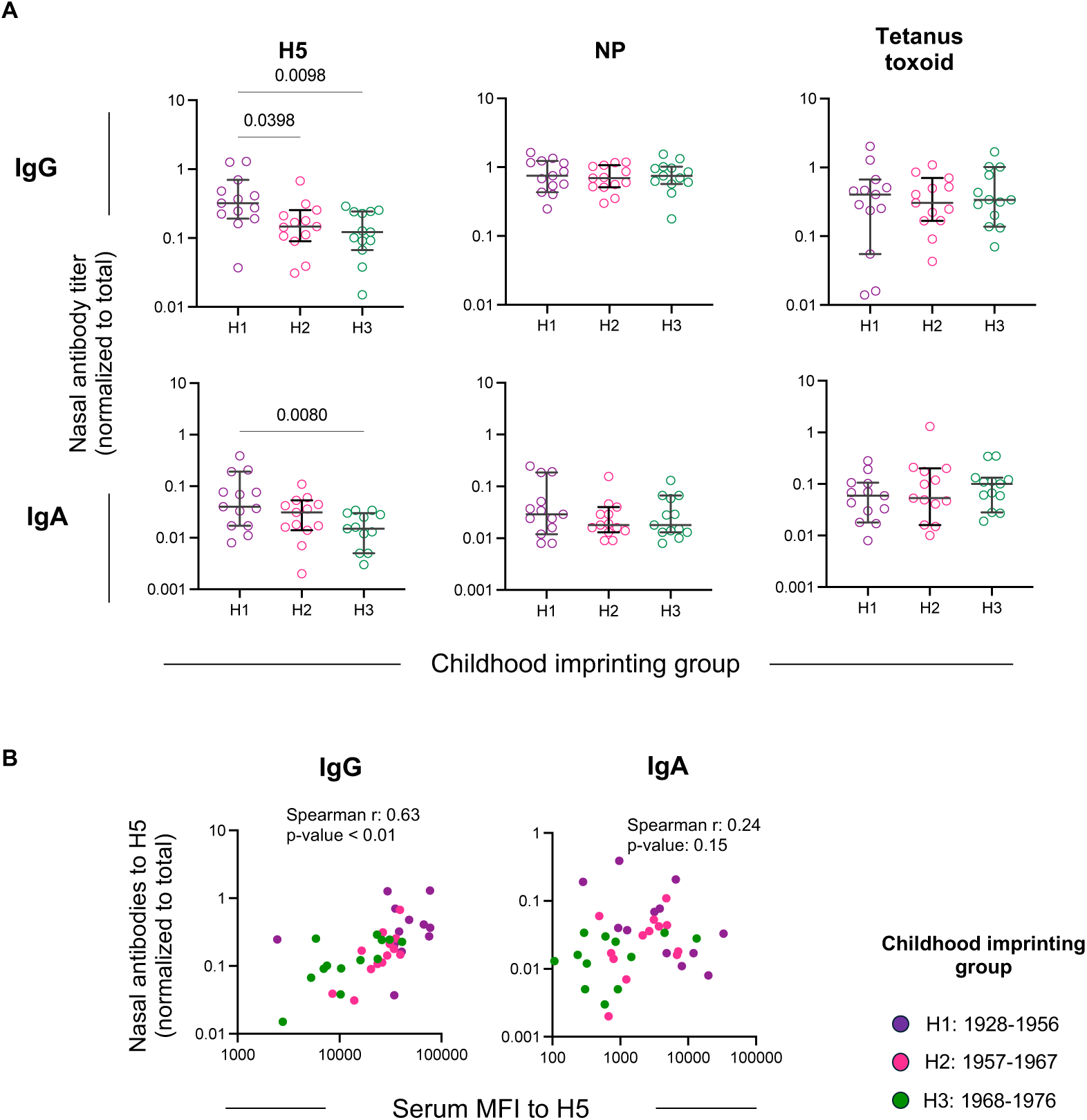
Pre-existing nasal antibodies against avian H5 IAV vary by birth year. **(A)** IgG and IgA antibodies specific for H5 (A/Fujian-Sanyuan/21099/2017), the IAV NP and tetanus toxoid were determined in nasal fluid using a multiplex bead assay. The levels of antigen-specific antibodies were normalized to the total IgG or IgA detected in each sample. Medians and 95% CIs (n=13/group) are shown, and p-values were determined by a Kruskal-Wallis test with Dunn’s multiple comparisons test and only comparisons with p-values <0.05 are shown. **(B)** Spearman correlations between antibody titers to H5 or NP between serum and nasal fluid (n=39).

## Discussion

The spread of 2.3.4.4b H5 viruses across a plethora of animal species highlights the need for preparedness should this high pathogenicity IAV acquire adaptations that support human-to-human transmission. An important aspect of pandemic preparedness is understanding pre-existing population immunity and heterogeneity in susceptibility. Here we show that pre-existing antibodies against the avian H5 can be detected in adults, but their levels vary according to early life exposure to group 1 or group 2 HAs. We demonstrate that this does not translate into differences in serum neutralization activity or differences in MBC pools, but it does result in differential levels of nasal IgG and IgA against H5. These findings have potential implications for the use of H5 vaccines as well as our understanding of immune imprinting.

The observation of varying levels of pre-existing antibodies to H5 by year of birth from our cohort in Australia is consistent with findings from independent studies from different geographic regions(12, 13), suggesting this is a generalisable feature of the global population. An important aspect of our analyses is the use of conserved control antigens like NP, that allow us to exclude the possibility of recent exposures to seasonal IAV, that could result in higher levels of antibodies, including towards distant antigens like H5(16, 17). The lower antibody levels detectable in younger individuals is consistent with the higher burden of zoonotic H5 infection in that age group(9). These findings collectively suggest that, in the event of an H5 IAV pandemic, those born after 1968 might be disproportionally impacted compared to individuals born prior to 1968. While this may imply increasing vaccine access to the those born after 1968, maintaining immunity in older individuals (those born before 1968) is often a priority, given their heightened susceptibility to severe influenza disease due to higher rates of comorbidities(24). Importantly, despite the association between ELISA IgG titres and protection from symptomatic seasonal influenza (7, 8, 25), there is a critical gap in the data required to translate the detected levels of IgG antibodies (in serum or nose) to protection curves, in order to quantify how these antibody measures directly translate to protection at the population level. In particular, given the associations of nasal antibodies with protection from influenza challenge(26) and with reduced viral shedding(27), an improved understanding of the protection afforded by nasal antibody titres against IAV infection and severe disease in humans would assist with further preparedness modelling that could inform appropriate public health measures in the event of a significant outbreak.

It is also important to consider if the differential levels of pre-existing antibodies may impact subsequent vaccine responses with H5 antigens. Recent studies suggest that post-vaccination antibody responses to H5 vaccines do not differ by birth cohort, at least in early timepoints post vaccination (day 28)(13). This might be explained by the similar frequencies of H5-specific MBCs detected in our cohort which provide an immediate source of cross-reactive antibodies. It remains important to determine if the development of *de novo* H5 head-specific antibody responses is impacted by higher levels of pre-existing stem immunity.

Our study supports the idea that most pre-existing antibodies detected in serum target the highly conserved stem region of group 1 HAs. Interestingly, the higher levels of HA stem-specific antibodies did not result in considerable neutralisation activity. The limited neutralisation activity is consistent with other studies using live IAV-based microneutralization assays(12, 16, 28). Studies using lentivirus-based pseudoviruses have reported more readily detectable neutralisation activity(29). This is likely due to a higher sensitivity of pseudovirus-based assays, but the degree to which such low neutralisation activity contributes to protection is unclear. Indeed, animal studies suggest that stem-specific antibodies primarily protect via Fc-mediated activity rather than direct virus neutralising(30, 31). While our study has not assessed the pre-existing levels of NA-specific antibodies, recent reports suggest considerable pre-existing immunity against N1, and N1-specific pre-existing immunity may contribute to neutralisation activity(32, 33). It is also important to note that pre-existing immunity to the group 2 H7 antigen was lower than that to H5, and H7-specific serum IgG did not vary by year of birth. This highlights a likely immunity gap towards group 2 HAs, that may suggest greater risk of these subtypes.

An important finding from our study is that the differential levels of pre-existing serum antibodies observed between birth cohorts is also reflected in nasal IgG and IgA levels. The levels of H5-specific antibodies detected in nasal fluid were strongly correlated with the levels detected in serum, and we therefore cannot conclude whether the antibodies detected in the nose are locally derived, transcytosed from serum or, most likely, a mixture of both. Delineating these possibilities would be important to understand how well the immunological biases detected in serum represent any immunological biases present in the mucosa.

We previously demonstrated that pre-existing immunity to future influenza B virus (IBV) variants varies according to childhood imprinting of different birth cohorts and is consistent with the differential susceptibility of different birth cohorts to the IBV antigenic lineages(34). The observation of differential pre-existing immunity against H5 according to childhood imprinting and in a manner consistent with epidemiological data further supports the link between immunological biases resulting from early life exposures and subsequent susceptibility later in life. This highlights the need for further studies to dissect heterogeneity in population immunity against antigenically variable pathogens. This could inform the integration of serological data in public health assessments to identify high-risk groups, as well as to refine pandemic models that account for baseline seroprevalence to accurately capture transmission dynamics and the potential impact of intervention strategies

## Materials & Methods

### Human serum samples

Samples from adults were collected under study protocols that were approved by the University of Melbourne Human Research Ethics Committee (2056689, 30807-59288, 21198-48733, 31386-63306).. All participants provided written informed consent in accordance with the Declaration of Helsinki. Peripheral blood mononuclear cells (PBMCs) were isolated via Ficoll-Paque separation, cryopreserved in 10% DMSO/FCS and stored in liquid nitrogen. Serum samples were collected and stored at -80° until use. Nasal fluid samples were collected via Nasosorption FX-I devices (Mucosal Diagnostics) and eluted in assay buffer (#AB-33K; Abacus Dx), before being stored in -80°C until use.

### Viruses and reagents

A recombinant IAV with the internal gene segments from the mouse adapted A/PR/8/1934 and the HA and NA from A/Astrakhan/3212/2020 (H5N8) was propagated in 10–12-day-old embryonated chicken eggs. Recombinant HA proteins (A/Sydney/5/2021 H1pdm09, A/Solomon Islands/03/2006 H1, A/Darwin/6/2021 H3, A/PR8 group 1 stabilised stem, A/Finland/486/2004 H3 group 2 stabilised stem, A/Shanghai/02/2013 H7, A/Canada/720/2005 H2, A/Fujian-Sanyuan/21099/2017 H5) were produced in-house in mammalian cells as previously described(18, 19, 35, 36). The antigenic integrity of these proteins was validated using cognate ferret antisera or monoclonal antibodies (CR9114, FR-1122). Recombinant IAV NP (#40947-V08B) and simian immunodeficiency virus glycoprotein gp120 (40415-V08H) were obtained from SinoBiological. Tetanus toxoid was obtained from Sigma Aldrich (#T3194-25ug).

### ELISA

Maxisorp 96-well plates (Thermo Fisher) were coated with 2 µg/mL recombinant HAs or NP overnight at 4°C. For stabilised stem antigens, plates were coated with 0.83µg/ml, which equimolar to 2 µg/mL of full-length HA. Plates were blocked with 200 µL of 1% fetal calf serum (FCS) in PBS for 1 hour, and then serially diluted sera were added and incubated for 2 hours at room temperature. Serum samples were prepared by making a serial of fourfold dilutions starting from 1:100 dilution. Plates were washed in PBS-T (0.05% Tween-20 in PBS) and PBS before incubation with a 1:20,000 dilution of HRP-conjugated rabbit anti-human IgG (LGC Clinical Diagnostics, Milford, USA) for 1 hour at room temperature. Plates were washed and developed using 100 µL 3,3’,5,5’-Tetramethylbenzidine (TMB) substrate and stopped by 50 µL 0.16M sulfuric acid, and read at 450 nm. Endpoint titres were determined using 2x the average signal of background (2° HRP antibody only) wells as a cut-off by a sigmoid 4PL curve in Graphpad Prism v9. Samples at the limits of detection were assigned the value of the highest or lowest dilution tested.

### HA-specific memory B cell staining

Surface staining of B cells within cryopreserved human PBMCs was performed as previously described(18–20, 35, 37). PBMCs were incubated with recombinant H5 [A/Fujian-Sanyuan/21099/2017(H5N6)] probes conjugated to streptavidin-APC or streptavidin-PE, as well as the following monoclonal antibodies: CD14-BV510 (M5E2, 1:200), CD3-BV510 (OKT3, 1:400), CD8a-BV510 (RPA-T8, 1:1000), CD16-BV510 (3G8, 1:300), CD10-BV510 (HI10a, 1:500) (all from BioLegend), unconjugated streptavidin-BV510 ( 1:400), IgG-BV786 (G18-145, 1:50), IgD-PeCy7 (IA62, 1:100) (all from BD Biosciences) and CD19-ECD (J3-119, 1:100, Beckman Coulter, Brea, USA). Probes and antibody staining was performed in PBS with 1% foetal calf serum (FCS). For all samples, cell viability was assessed using Aqua Live/Dead amine-reactive dye (Thermo Fisher). Between 1.5-4.5 million events were collected on an LSR Fortessa (BD). Analysis was performed using the FlowJo software version 10.10 (BD). H5-specific B cells were identified using the gating strategy is Figure S1.

### Microneutralization assay

The neutralization activity of serum was examined in a microneutralization assay using a recombinant IAV with the internal gene segments from the mouse adapted A/PR/8/1934 and the HA and NA from A/Astrakhan/3212/2020 (H5N8), with the multi-basic cleavage site removed from the HA. Briefly, MDCK cells were seeded in 96-well plates at 3 × 10^4^ per well 1 day before the assay and incubated at 37 °C overnight. Serum samples were heat inactivated at 56 °C for 30 min and serially diluted (two-fold, starting at 1:10) in cell media (high glucose and sodium pyruvate DMEM supplemented with GlutaMAX, MEM non-essential amino acids, sodium bicarbonate, HEPES, Penicillin Streptomycin and Amphotericin B). Serum samples were incubated for 60 min at 37 °C with an equivalent volume of 100 TCID50 units of virus diluted in VIM. After incubation, MDCK cells were washed twice in PBS. VIM (100 μl) supplemented with 2 μg ml^−1^ TPCK-treated trypsin and 100 μl virus–serum mix were added to the MDCK monolayer. Control wells of virus alone and VIM alone were included on each plate. Virus input was back titrated on MDCK cells. Cells were incubated for 3 days at 37 °C, and the presence of virus was determined by haemagglutination assay. Briefly, 25 μl of supernatant from each well was mixed with 25 μl of 1% (v/v) turkey erythrocytes and incubated for 30 min at room temperature, and the presence of virus was recorded. Microneutralization titres were determined using the reciprocal of the highest dilution at which virus neutralization was observed.

### Bead-based multiplex array

A custom multiplex bead-based array consisting of recombinant H5 and IAV NP was used to assess influenza-specific IgG and IgA binding in serum and nasal fluid, as previously described(38). Tetanus toxoid was included as a positive control and simian immunodeficiency virus glycoprotein gp120 was included as a negative control. Total IgG and IgA antibodies in serum and nasal fluid were measured using beads coupled to anti-human IgG and IgA respectively (MT145; MT57; Mabtech). Briefly, capture-antibody or antigen-coupled beads were first incubated with serum or nasal samples overnight at 4°C, washed and then incubated with the respective biotinylated detectors (anti-human IgG MT78; anti-human IgA MT20; Mabtech) for 2 hours at room temperature. After washing, beads were incubated with streptavidin-R-phycoerythrin (Thermo Fisher Scientific) for 2 hours at room temperature. Finally, beads were washed and read on the Intelliflex (Luminex). Assays were performed in duplicate. The antigen-specific antibody binding titres were normalized to the total isotype (IgG or IgA) content in each nasal sample.

### Statistical analysis

For the initial cohort of n=144, mean endpoint titres (log scale) with 95% confidence intervals (CI) for each imprinting group were calculated from a linear regression model, using the emmeans (version 2.0.0)(39) package in R (version 4.4.0)(40). Separate models were run for each antibody type. For each model, pairwise contrasts between birth cohorts were estimated, and p-values were adjusted using Tukey’s method to account for multiple comparisons across the four birth cohorts.

Data visualisation of endpoint titres against year of birth was performed using ggplot2 package (version 4.0.2)(41). Pairwise Pearson correlations were performed for each combination of antibodies against HAs from IAV subtypes and stem antigens. No multiple testing adjustments were made. All visualisation and correlation analyses were performed using R (version 4.4.0).

For the second cohort (n=39) data visualisation was performed in GraphPad Prism v9. Normality was assessed using the D’Agostino & Pearson test. Groups were compared using an ANOVA with Tukey’s correction for multiple comparisons for normally distributed data, or a Kruskal-Wallis test with Dunn’s multiple comparisons test. Multiple comparisons were accounted for within each analysis, where pairwise comparisons were made between each birth cohort. Spearman rank correlations were performed.

## Data and Code availability

The codes used for analyses and figures 1 and 2 are available via 10.5281/zenodo.20060813

## Acknowledgements

The work has been generously supported by the Morningside Foundation, by the Australian National Health and Medical Research Council, by a Doherty Institute Collaborative Seed Grant and by the Cumming Global Centre for Pandemic Therapeutics. The WHOCCRRI is supported by the Australian Government Department of Health. The funders had no role in study design, data collection and analysis, decision to publish or preparation of the paper. For the purposes of open access, the author has applied a CC BY public copyright licence to any Author Accepted Manuscript version arising from this submission.

## Author contributions

M.K. conceived the study. M.K., K.J.S., E.R. and M.A. performed experiments. I.B., J.A., H.K., J.A.J., A.K.W., A.W.C. and S.J.K. provided samples and/or critical reagents. M.K., K.J.S., P.E., and D.J.P. analysed data. M.K., K.J.S., P.E., and D.J.P. drafted the manuscript. All authors read and approved of the manuscript.

## Supplementary figures

**Figure S1.**
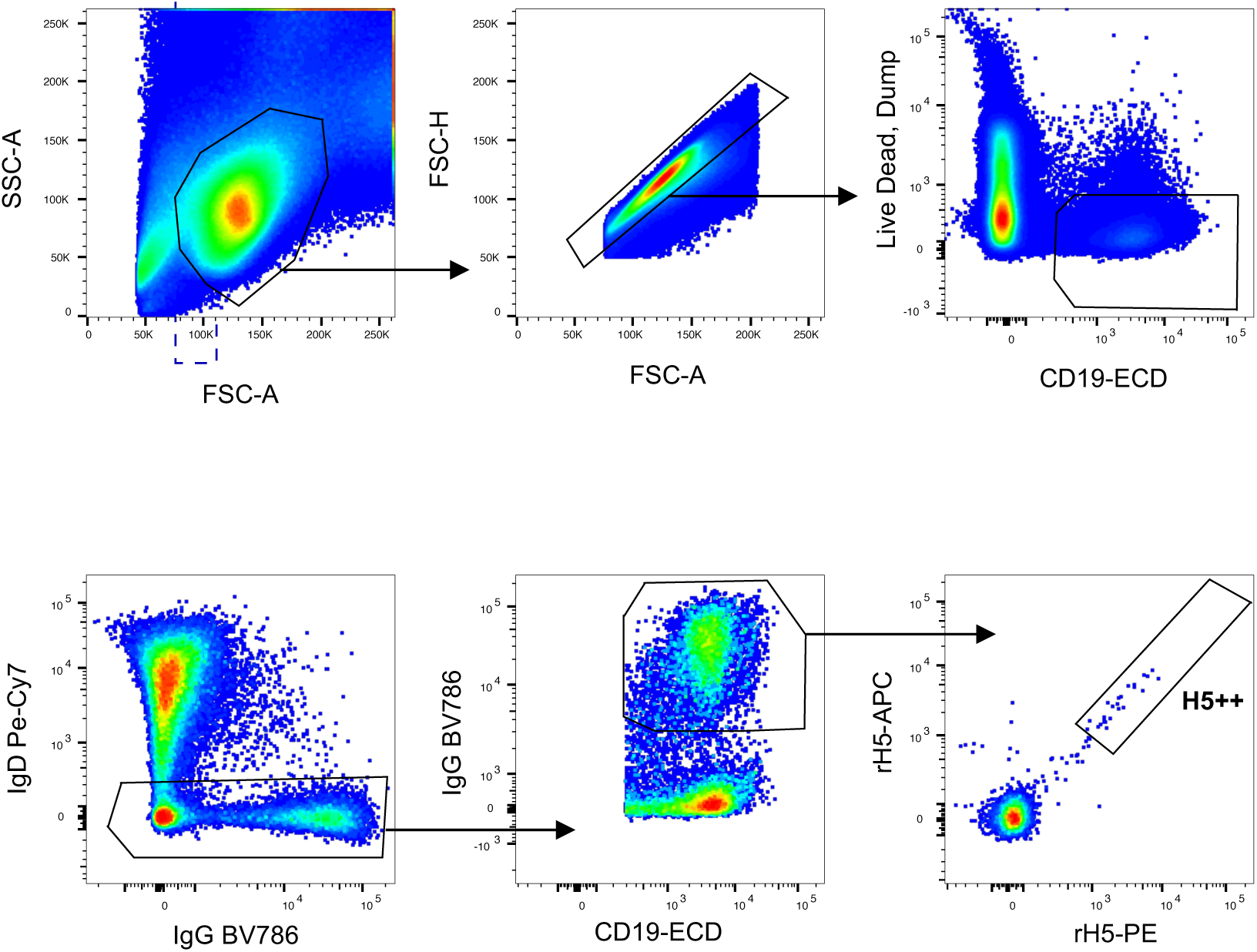
Gating strategy for the identification of H5 specific memory B cells. A/Fujian-Sanyuan/21099/2017 H5-specific B cells are shown after gating on IgG^+^IgD^-^CD19^+^CD3^-^CD8^-^CD16^-^CD14^-^CD10^-^ live single lymphocytes.

**Figure S2.**
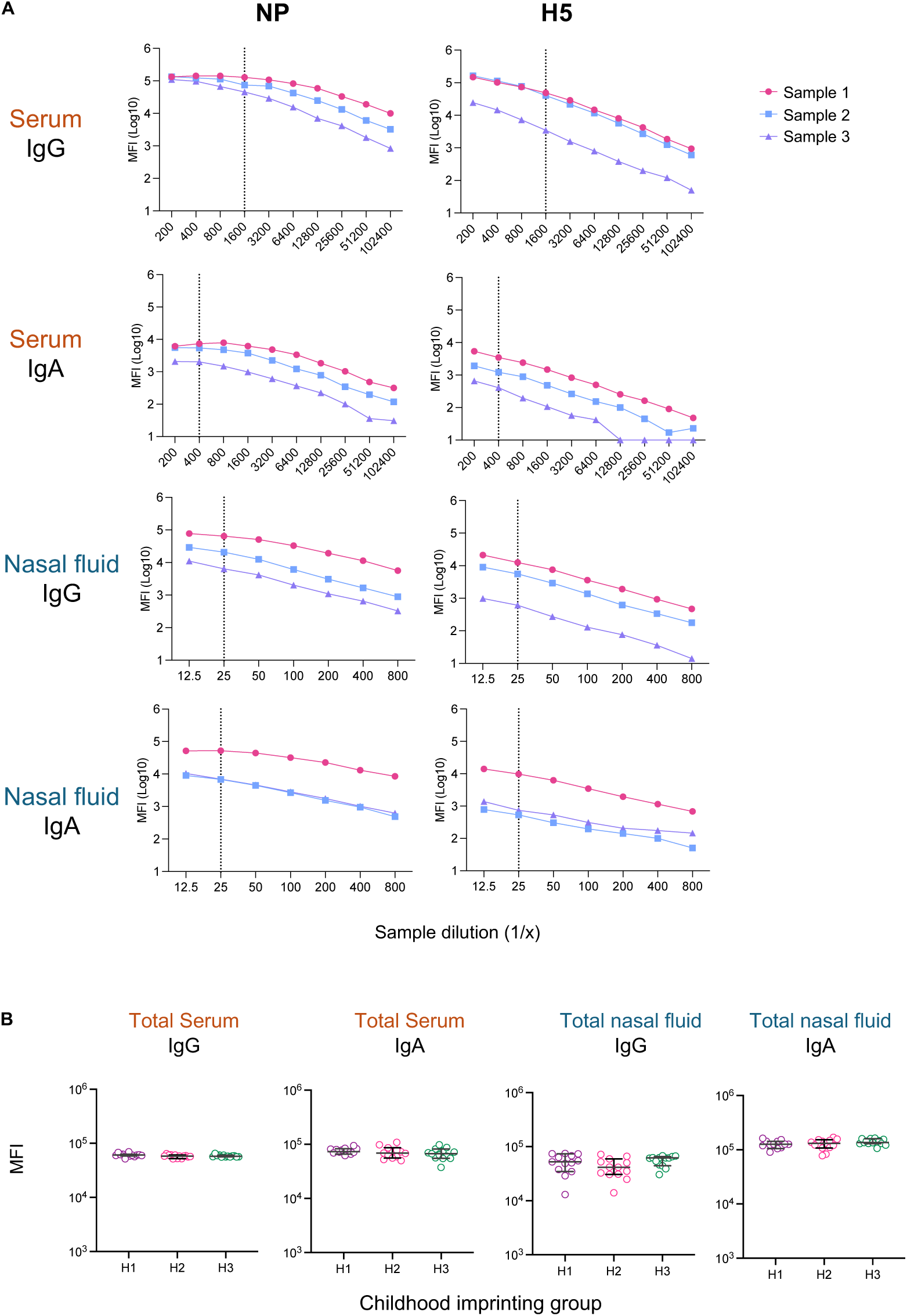
Multiplex bead assay set up. **(A)** IgG and IgA titration curves for IAV NP and H5 coupled beads with serum and nasal fluid from 3 representative samples. Dotted lines depict the working dilutions chosen for each sample type and antibody isotype respectively. **(B)** Total levels of IgG and IgA detected in each sample. Medians and 95% CIs (n=13/group) are shown, and p-values were determined by a Kruskal-Wallis test with Dunn’s multiple comparisons test and were all >0.05.

**Figure S3.**
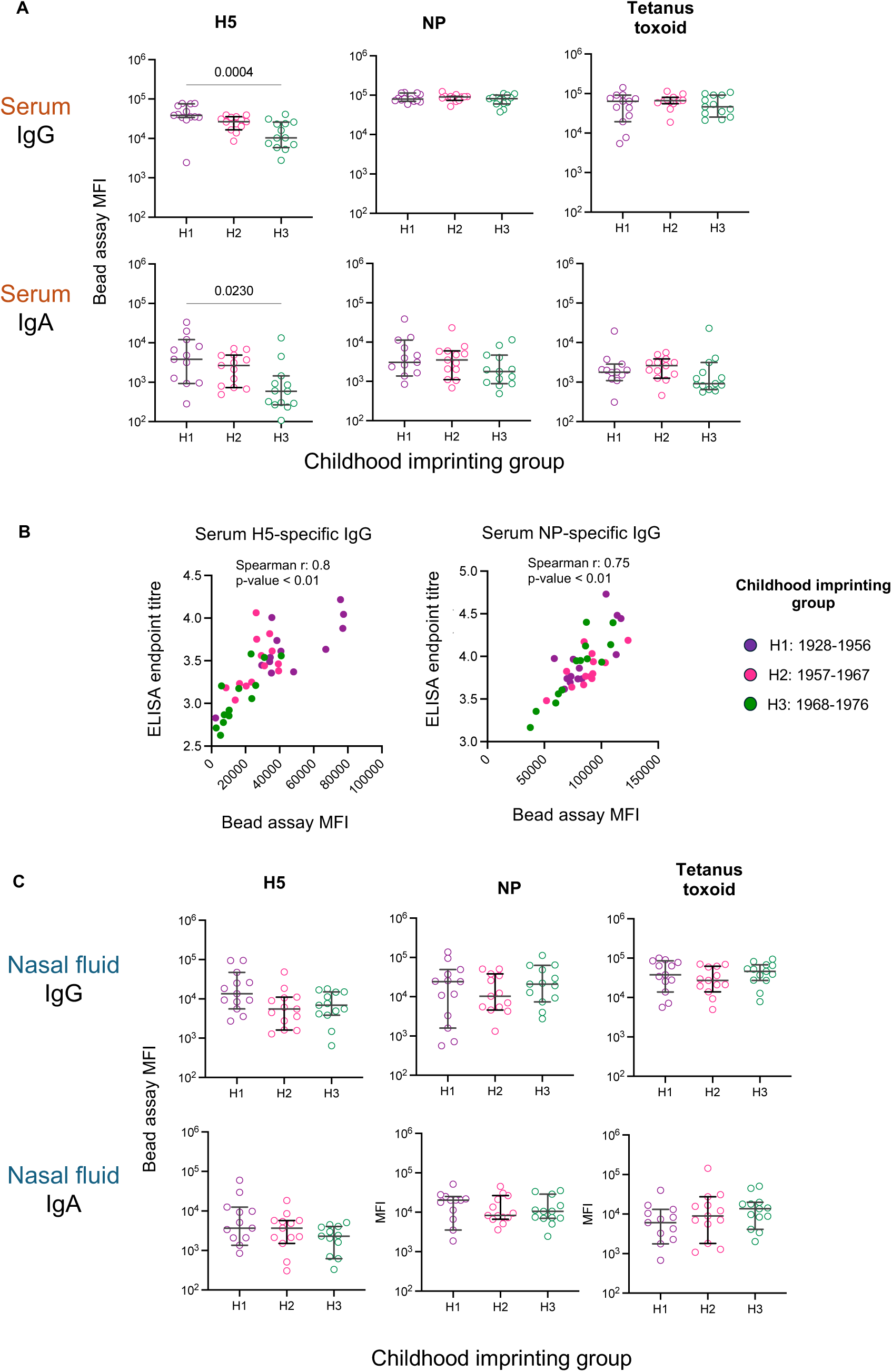
Serum and nasal fluid IgG and IgA against H5 and control antigens. **(A)** IgG and IgA antibodies specific for H5 (A/Fujian-Sanyuan/21099/2017), the IAV NP and tetanus toxoid were determined in serum using a multiplex bead assay. Medians and 95% CIs (n=13/group) are shown, and p-values were determined by a Kruskal-Wallis test with Dunn’s multiple comparisons test and only comparisons with p-values <0.05 are shown. **(B)** Correlation between antibody titers to H5 or NP determined by ELISA (Figure 2) or by the bead multiplex assay (n=39). **(C)** IgG and IgA antibodies specific for H5 (A/Fujian-Sanyuan/21099/2017), the IAV NP and tetanus toxoid were determined in nasal fluid using a multiplex bead assay. Medians and 95%CI (n=13/group) are shown, and p-values were determined by a Kruskal-Wallis test with Dunn’s multiple comparisons test and were all >0.05.

